# High capacity of Brazilian *Aedes aegypti* populations to transmit a locally circulating lineage of Chikungunya virus

**DOI:** 10.1101/2023.10.23.563517

**Authors:** Amanda Cupertino de Freitas, Fernanda Oliveira Rezende, Silvana Faria de Mendonça, Lívia Vieira Baldon, Emanuele Guimarães Silva, Flávia Viana Ferreira, João Paulo de Almeida, Siad Cedric Amadou, Bruno Almeida Marçal, Sara Grangeiro Comini, Marcele Neves Rocha, Hegger Machado Fritsch, Marta Giovanetti, Luiz Carlos Alcantara, Luciano Andrade Moreira, Alvaro Gil Ferreira

## Abstract

The global incidence of chikungunya has surged in recent decades, with South America, particularly Brazil, experiencing devastating outbreaks. The primary vector for transmitting CHIKV in urban areas is the mosquito species *Aedes aegypti*, which is very abundant in Brazil. However, little is known about the impact of locally circulating CHIKV genotypes and specific combinations of mosquito populations on vector competence. In this study, we analyzed and compared the infectivity and transmissibility of a recently isolated CHIKV-ECSA lineage from Brazil among four *Ae. aegypti* populations collected from different regions of the country. When exposed to CHIKV-infected mice for blood feeding, all mosquito populations showed high infection rates and dissemination efficiency. Moreover, using a mouse model to assess transmission rates in a manner that better mirrors natural cycles, we observed that these populations exhibit highly efficient transmission rates of CHIKV-ECSA. Our findings underscore the robust capability of Brazilian *Ae. aegypti* populations to transmit the locally circulating CHIKV-ECSA lineage, potentially explaining its higher prevalence compared to the Asian lineage also introduced in Brazil.

**Author Summary:** Chikungunya virus (CHIKV) is an alphavirus transmitted by mosquitoes, gaining attention due to its rapid global spread and public health impact. Initially isolated in Tanzania in 1952, it was confined to Africa, Asia, and the Indian subcontinent. Local transmission of CHIKV in the Americas began in December 2013, leading to a 2014 epidemic. The *Aedes aegypti* mosquito, prevalent in Brazil, is suspected as the primary vector. Yet, our understanding of how different mosquito populations and virus lineages impact spread is limited. In this Brazilian study, we collected mosquitoes from four regions, exposing them to the local African ECSA lineage of CHIKV. Remarkably, all mosquito populations exhibited high infection rates and efficient virus transmission to vertebrate hosts. This research sheds light on why the ECSA chikungunya lineage is spreading rapidly in Brazil. The Brazilian *Ae. aegypti* mosquitoes appear to possess exceptional capabilities in transmitting African ECSA lineage of CHIKV, potentially contributing to its rapid dissemination within the country and neighboring regions. Comprehending these dynamics is vital for developing strategies to control and mitigate the impact of chikungunya in affected areas.

## Introduction

Chikungunya virus (CHIKV) is a major threat to global public health, and over the last three decades, it has been responsible for epidemics that are increasing in frequency and geographic scale [1–3]. Currently, CHIKV circulates throughout the tropical and subtropical regions of Africa, South Asia, and South America[4–6]. CHIKV is an arthropod-borne virus (arbovirus) that is transmitted mainly by Aedes mosquito species, and the infection commonly results in a febrile illness called chikungunya fever (CHIKF), which is usually self-limiting [1,3,7]. The main symptom of CHIKF is chronic and severe joint pain, which can be accompanied by an itchy maculo-papular skin rash and can lead to prolonged periods of functional disability. Severe complications, such as encephalitis and fatal outcomes may occur in individuals with comorbidities [3].

CHIKV is an alphavirus (Togaviridae family) that has a positive sense RNA genome encoding four non-structural proteins (nsP1-4) and five structural proteins (C, E3, E2, 6K, and E1) expressed from a subgenomic RNA [8]. Historically, CHIKV originated in sub-Saharan Africa where a transition occurred from an ancestral enzootic sylvatic cycle involving arboreal mosquitoes and nonhuman primates to an urban cycle where peri-domestic mosquitoes transmit the virus among humans [7]. According to phylogenetic studies, CHIKV first emerged in the urban cycle that encompasses humans in the eastern region of Africa before spreading to other parts of the world [4,6]. The virus is currently classified into three genotypes: West African, East-Central-South-African (ECSA), and Asian. More recently, the ECSA genotype has given rise to the Indian Ocean lineage (IOL), which has spread globally and caused numerous outbreaks since 2005 [2,9,10].

The distribution of CHIKV has rapidly expanded since its spread from Africa. The virus is currently being transmitted locally in more than 100 countries across various regions, including the Americas, the Caribbean, North America, the Western Pacific, Southern Europe, Southeast Asia, and Oceania [11–21]. Due to its geographical spread and ability to cause incapacitating disease, CHIKV was included by the World Health Organization (WHO) in the Priority Blueprint under the Global Arbovirus Initiative (GAI), which was launched in 2022 to increase global awareness of the potential risk of epidemics[22]. Indeed, the number of outbreaks of CHIKV has increased dramatically in recent years, especially in the South American region. Currently, Brazil has the highest number of chikungunya cases across the entire region [23,24].

There are currently two confirmed CHIKV lineages circulating in Brazil: the Asian lineage and the African ECSA lineage, both introduced in 2014 [9,24–26]. Nonetheless, it appears that the ECSA lineage is the primary genotype currently in circulation in the country [10,23,26–28]. Although the epidemiology of CHIKV depends on several complex factors, including host and viral factors, vector factors such as intrinsic infection ability and transmission efficiency play an important role in the dynamics and establishment of different lineages. In Brazil, the main vectors associated with the transmission of these CHIKV lineages are the mosquitoes *Aedes aegypti* and *Aedes albopictus*, both of which are widely distributed across the country [14,18,20,29]. Previous studies have shown that these two Aedes species are competent vectors for both the ECSA and Asian lineages across different world regions [15,29–36]. However, in Brazil, the impact of locally circulating CHIKV genotypes and specific combinations of local *Aedes aegypti* populations on vector competence has not been thoroughly examined. Here, we evaluated and compared the infectivity and transmissibility of a CHIKV-ECSA lineage recently isolated in Niterói, Brazil [10,24], among four *Ae. aegypti* populations collected from different regions of the country. We observed that all *Ae. aegypti* populations displayed high infection rates and dissemination efficiency when exposed to CHIKV-infected AG129 mice for blood feeding. Additionally, we observed that all populations were highly efficient in transmitting CHIKV to a vertebrate host (naïve AG129 mice) as early as eight days post-infection, indicating that the circulating CHIKV-ECSA lineage in Brazil is readily transmitted by local *Ae. aegypti* populations.

## Methods

### Mosquito Collection and Establishment of the Populations

Mosquitoes were sampled in four sites in Brazil in four different states: Araraquara city in São Paulo state (ARA); Jaboticatubas in Minas Gerais state (JAB), Petrolina city in Pernambuco state (PET) and in Porto Alegre city in Rio Grande do Sul during the months January and February of the year 2022. Eggs were collected and shipped to the Laboratório of Mosquitos Vetores (MV) at the Instituto René Ra-chou-Fiocruz Minas, Belo Horizonte, Brazil. All mosquitoes were reared under insectary-controlled conditions, 28°C and 70–80% relative humidity, in a 12/12 h light/dark cycle. Eggs were placed in plastic trays containing two liters of filtered tap water, supplemented with fish food (Tetramin, Tetra tablets) for hatching, and larvae were maintained at a density of 200 larvae per tray. After emerging, adults were kept in 30 cm × 30 cm × 30 cm BugDorm insect cages, where mosquitoes were fed with 10% sucrose solution ad libitum. For the establishment of each laboratory population 100 females and 100 males were used.

### Virus Strain, Viral Propagation and Titration

To evaluate the mosquito vector competence, we used CHIKV-ECSA strain isolated from a human patient in Niterói, Rio de Janeiro state, Brazil, in 2018. The Serum sample from the patient with CHIKV-like symptoms was used for molecular diagnostics. After molecular screening, nucleic acid extraction and purification were performed following the manufacturer’s recommendations, using the Magmax Pathogen RNA/DNA and KingFisher Plex Purification System (ThermoFisher) kits. RT-qPCR for CHIKV RNA detection followed an adapted protocol from Fritsch and coauthors[24]. This CHIKV lineage was propagated in C6/36 *Ae. albopictus* cells. C6/36 cells were maintained on L15 medium, supplemented with 10% FBS (fetal bovine serum) and 1× Antibiotic Antimycotic (Gibco), as described [37]. Cells were seeded to 70% confluence and infected at a multiplicity of infection (MOI) of 0.01 and maintained for three to five days at 28°C. The supernatant was collected and clarified by centrifugation to generate virus stocks that were kept at −80°C prior to use. Mock supernatants used as controls were prepared under the same procedure, without virus infection. Titration was performed in Vero cells using the plaque assay method to determine the viral titer. We allowed the virus to adsorb for 1 h at 37°C, then an overlay of 2% carboxymethyl cellulose (CMC) in DMEM with 2% FBS was added. Plates were incubated at 37°C and 5% CO2 for 3-5 days. Then, formaldehyde was added, and the cells were covered with a crystal violet stain (70% water, 30% methanol, and 0.25% crystal violet) to visualize the plaques.

### Mice Inoculation with CHIKV

CHIKV inoculation of AG129 mice (IFN α/β/γ R−/−) was accomplished by intra-peritoneal injection (IP) in four-week-old animals. We inoculated 10^5^ p.f.u. per animal. Following inoculation, the mice were visually monitored daily and scored for morbidity and mortality. For all experiments, the mice were bred and kept during the inoculation experiments in a specific-pathogen-free facility at Instituto René Rachou. The mice were maintained in a temperature- and humidity-controlled facility on a 12-hour light/dark cycle with food and water ad libitum. AG129 mice were bred and maintained at the Animal Facility of the Instituto René Rachou, Fiocruz Minas. Experiments were approved by the Institutional Animal Care and Use Committee, Comissão de Ética no Uso de Animais da Fiocruz (CEUA) and performed according to institutional guidelines (license number LW-26-20).

### Mosquito Infection with CHIKV

Five to seven-day-old mosquito females were transferred into cylindrical containers fitted with nylon mesh (0.88 mm hole size) on top and starved through sugar-deprivation for 24 h prior to mice blood feeding. All mosquito infections were performed using AG129 mice. Infected AG129 mice were anesthetized two days post infection, using ketamine/xylazine (80/8 mg per Kg^−1^). Subsequently, anesthetized mice were placed on the top of the netting-covered containers with mosquito females. Unshaved mice were placed in a prone position, with the entire ventral surface and limbs available to the mosquitoes. Mosquito females were allowed to feed on mice for 30 min. After blood feeding, fully engorged females were selected. All engorged females were placed in a container covered with nylon mesh with a cotton pad soaked with 10% glucose solution and with a plastic cup with soaked paper on the bottom for egg laying. Mice fed mosquito females were harvested individually for tissue dissection and subsequent RNA extraction at four- and eight-days post feeding. All mosquito infection experiments were conducted in an insectary with Biosafety Level 2 (BSL-2) with certification number 157/02 from the CTNBio (Comissão Técnica Nacional de Biossegurança).

### CHIKV Transmission from Mosquitoes to AG129 Mice

To evaluate the transmission efficiency of CHIKV in mosquito populations, we first fed five-to seven-day-old female mosquitoes on viremic AG129 mice. Eight days after the infection, female mosquitoes were allowed to feed on two to three-week-old anesthetized naive AG129 mice. One female mosquito was exposed to each AG129 mouse for 30 min, and after blood feeding, fully engorged females were selected and harvested individually for RNA extraction for arbovirus quantification. Three days after the AG129 mice were exposed to the mosquitoes, blood was collected for arbovirus quantification by RT-qPCR.

### RNA Extraction and RT-qPCR

RNA extraction from serum samples or mosquito samples was performed using the Trizol method reagent (Invitrogen, Carlsbad, CA, USA), as previously described [38]. Total RNA was extracted, according to the manufacturer’s instructions, with some modifications. 50 µL of serum or mosquito tissues were placed in a 1.5 mL Eppendorf tube, and then 200 µL of Trizol and two glass beads were added, then the samples were grounded. The total RNA extracted was reverse transcribed using M-MLV reverse transcriptase (Promega, Madi-son, WI, USA) and using random primers for initiation. Negative controls were pre-pared following the same protocol, without adding the reverse transcriptase. All real-time PCR reactions were performed using the QuantStudio 12K Real-Time PCR System (Applied Biosystems, Foster City, CA, USA), and the amplifications were performed using the SYBR Green PCR Master Mix (Applied Biosystems—Life Technologies, Foster City, CA, USA). The final reaction volume was 10 µL. The thermal cy-cling conditions were composed of Hold Stage (fast ramp to 95 ◦C, hold 20 s); PCR Stage (40 cycles of 95 ◦C, hold 15 s, fast ramp to 60 ◦C, hold 60 s); and Melt Stage (fast ramp to 95 ◦C, hold 15 s, fast ramp to 60 ◦C, hold 1 min, slow ramp of 0.05 ◦C/s to 95 ◦C, hold 15 s). All real-time PCR reactions were carried out in triplicate. The relative quantification of gene expression was determined using the 2ΔCt method, as previously described [39]. The viral RNA load was expressed relative to the endogenous control housekeeping gene, RPL32, for *Ae. aegypti* and RPL32 AG129 mice. For *Ae. aegypti*, the RPL32 primers were: Forward: 5’-AGC CGC GTG TTG TAC TCT G-3’ and Reverse: 5’-ACTTCT TCG TCC GCT TCT TG-3’. For mice, the RPL32 primers were: Forward: 5’-GCTGCC ATC TGT TTT ACG G-3’ and Reverse: 5’-TGA CTG GTG CCT GAT GAA CT-3’. For CHIKV, the primers were: Forward: 5’-AAG CTY CGC GTC CTT TAC CAA G-3’ and Reverse: 50-CCA AATTGT CCY GGT CTT CCT-3’.

### Statistical Analyses

Statistical analyses were conducted using the software R v4.2.3 (www.r-project.org). Logistic regression analyses were performed to assess the association between predictor variable (the different populations of *Aedes aegypti*) and the binary outcome variable (infection status) using a Generalized Linear Model (GLM) with family argument set to “binomial” and the link function to “logit” followed likelihood-ratio χ2 tests. To compare the differences in viral loads in the midgut and carcass samples among the different mosquito populations, we performed a Kruskal-Wallis test, followed by a post-hoc test (Dunn test), to determine the pairwise differences between the groups. *p*-values less than 0.05 were considered statistically significant [37].

## Results

### Collection and establishment of mosquito populations

To assess the vector competence of Brazilian *Ae. aegypti* mosquito populations for CHIKV, we analysed the susceptibility to infection, the ability of the virus to disseminate out of the midgut and the potential of being transmitted to a vertebrate host. For that we used a CHIKV ECSA strain recently isolated in Brazil (Table 1).

**Table 1.** Information of the Chikungunya strain used in this study.

| Virus | Viral Lineage | GenBank | Origin | Location | Isolation Year |
| --- | --- | --- | --- | --- | --- |
| CHIKV | ECSA | MK244647 | Human patient | Niterói, Brazil | 2018 |

These analyses were performed for four different *Ae. aegypti* populations covering four different Brazil states: Araraquara city in São Paulo state (ARA), Jaboticatubas city in Minas Gerais state (JAB), Petrolina state in Pernambuco state (PET) and Porto Alegre city in Rio Grande do Sul state (POA), (Fig 1 and Table 2). For that, we exposed five to seven-day-old female mosquitoes from all four populations to the same CHIKV-infected AG129 mice, a double-knockout strain lacking receptors for both type I (α, β) and type II (γ) interferons (IFNAR−/− IFNGR−/−DKO), for blood-feeding for 30 minutes. At four- and eight-days post-feeding (d.p.f.), we collected the mosquitoes and analysed the presence of CHIKV in the midgut and carcass to evaluate the infection rates and dissemination efficiency, respectively. Additionally, at eight days post-feeding, another set of mosquitoes was individually exposed to naïve AG129 mice to evaluate transmission efficiency.

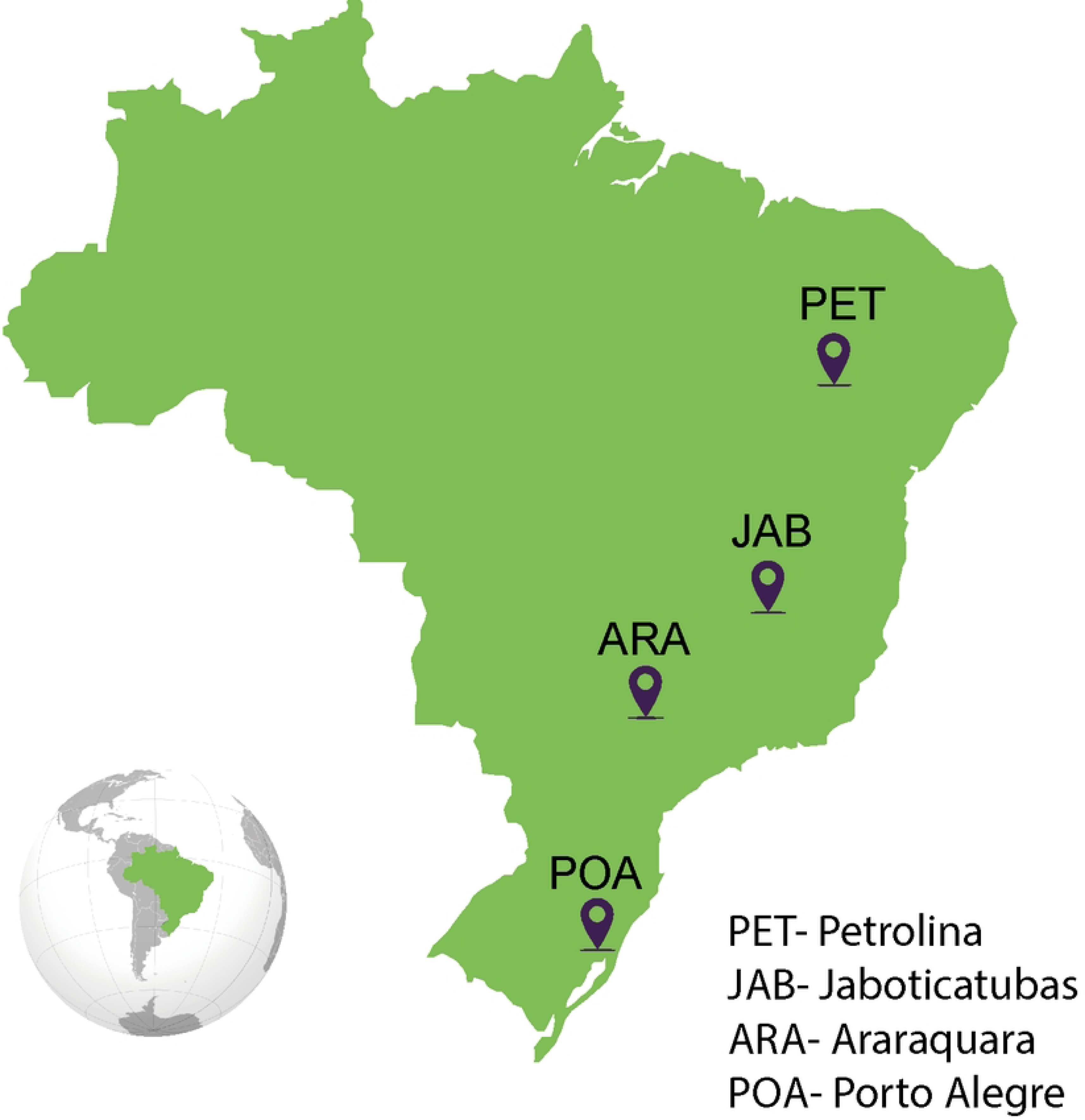

**Table 2.**
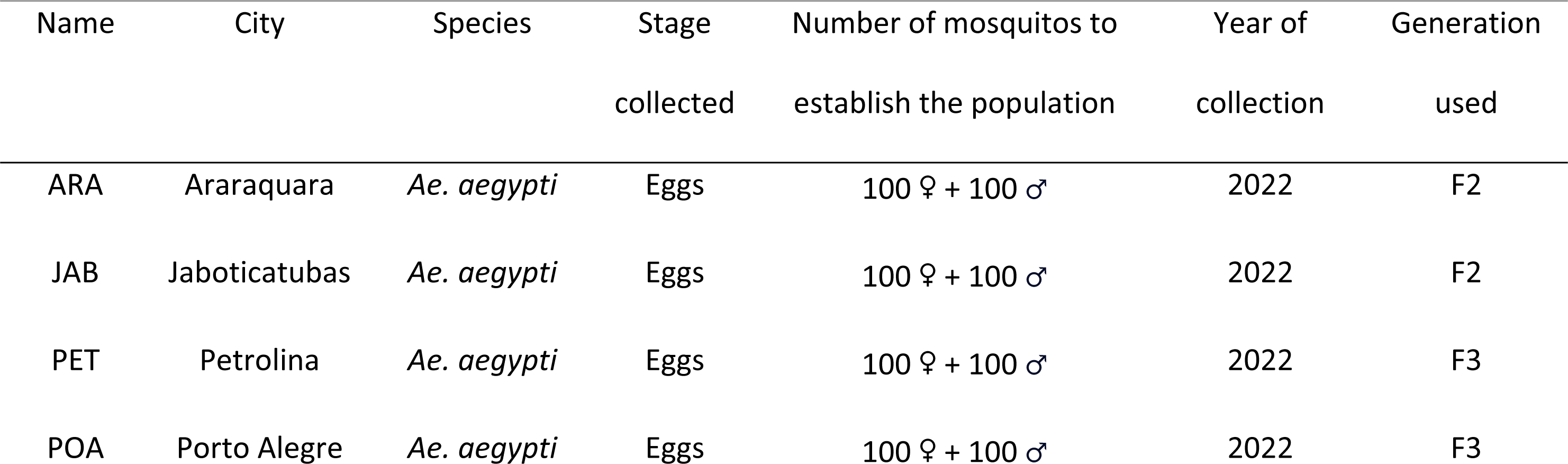
*Aedes aegypti* populations collected in Brazil.

| Name | City | Species | Stage<br>collected | Number of mosquitos to<br>establish the population | Year of<br>collection | Generation<br>used |
| --- | --- | --- | --- | --- | --- | --- |
| ARA | Araraquara | *Ae. aegypti* | Eggs | 100 ♀ + 100 ♂ | 2022 | F2 |
| JAB | Jaboticatubas | *Ae. aegypti* | Eggs | 100 ♀ + 100 ♂ | 2022 | F2 |
| PET | Petrolina | *Ae. aegypti* | Eggs | 100 ♀ + 100 ♂ | 2022 | F3 |
| POA | Porto Alegre | *Ae. aegypti* | Eggs | 100 ♀ + 100 ♂ | 2022 | F3 |

### Different populations of *Aedes aegypti* across Brazil exhibit high and similar CHIKV infection rates as well dissemination efficiency

Using the locally isolated CHIKV-ECSA strain, we assessed the infection rates, which were calculated as the proportion of blood-fed mosquitoes that tested positive for the virus in the midgut relative to the total number of blood-fed mosquitoes, in four *Ae. aegypti* populations (Fig 2.A). We found that infection rates were consistently high and uniform across all four Brazilian populations tested, starting as early as 4 days post feeding, and continuing through to 8 days post feeding (Fig 2.B, D). Ranging from 82% at 8 d.p.f for POA population to 100% at 4 d.p.f. for JAB population. Using a logistic regression model to estimate the relationship between the infection rates and the different populations, we found no significant differences between the four mosquito populations for both time points 4 d.p.f (*p*-value = 0.2093, S1 Table) and 8 d.p.f. (*p*-value = 0.5197, S1 table). Even when we analysed the data from both time points together, we found no significant differences between the populations (*p*-value = 0.7393, S3 Table). As shown in Fig 2.C, midgut viral loads at 4 d.p.f. were similar between the four mosquito populations tested (Kruskal-Wallis test). However, at 8 d.p.f., we observed that the PET population exhibited significantly higher amounts of CHIKV in the midgut compared to the other three populations tested (PET-ARA *p*-value = 0.00465; PET-JAB *p*-value = 0.000156; PET-POA *p*-value < 0.0001, Kruskal-Wallis test, Fig 2.E).

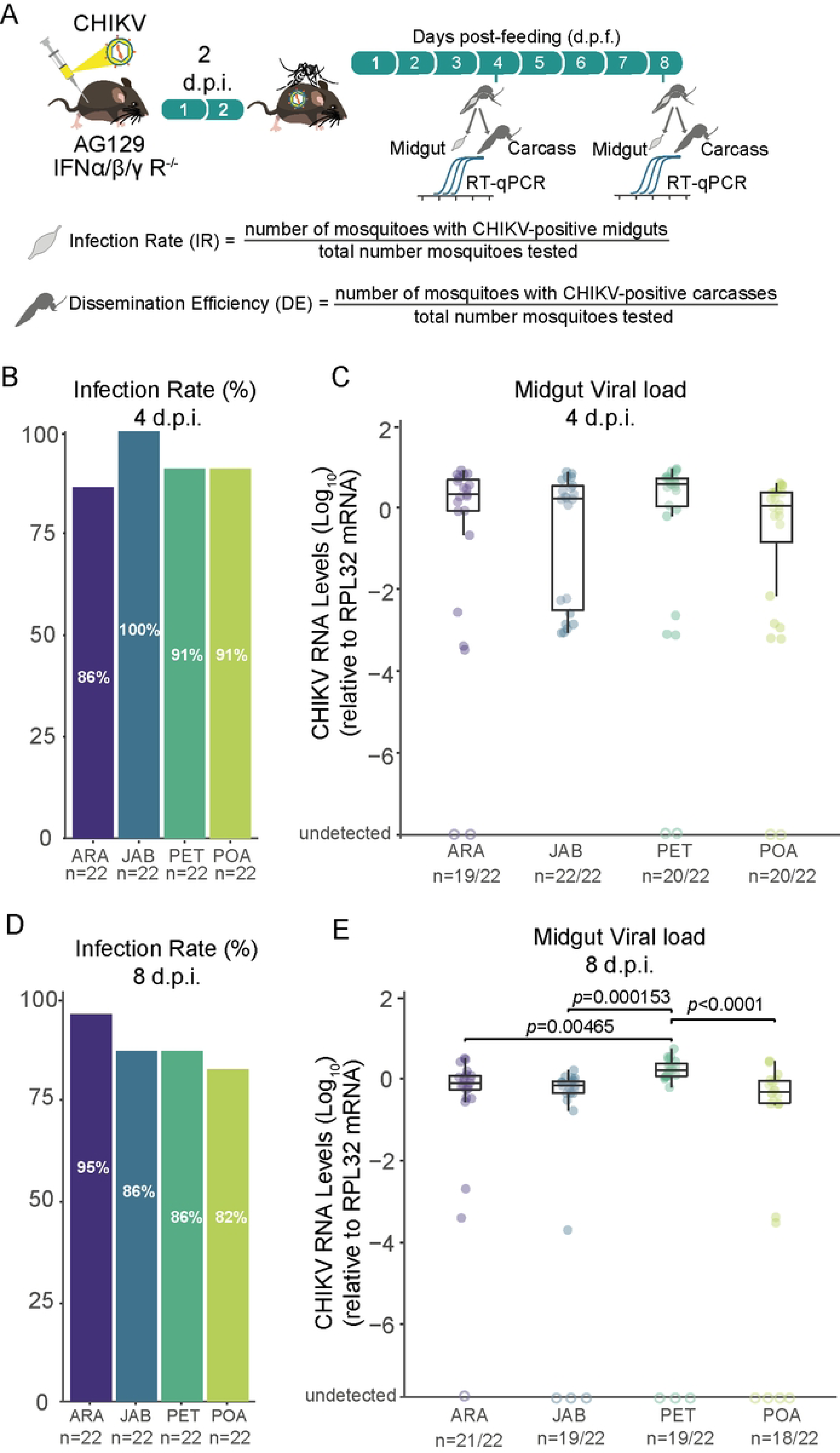

### CHIKV show high dissemination efficiency mosquitoes of Brazilian *Aedes aegypti* populations

To compare how well CHIKV spreads from the mosquito midgut to other body parts in the different *Ae. aegypti* populations, we measured the percentage of blood-fed mosquitoes with the virus in their carcass (their bodies excluding the midgut) out of the total number of blood-fed mosquitoes from four Brazilian populations. Notably, we observed that CHIKV was already present in the carcass as early as 4 d.p.f. in most of the mosquitoes tested (Fig 3.A). For both the 4 and 8 days d.p.f. time points, we observed consistently high dissemination rates across all four populations tested, ranging from 95% for the Araraquara population at 4 days post feeding to 80% for the Porto Alegre mosquitoes at 8 days post feeding (Fig 3.A, C). Nevertheless, no significant differences were found between the populations for either the 4 or 8 days d.p.f. time points (*p*-value = 0.012 and *p*-value = 0.6602, respectively; see S2 Table). We observed some variation in the viral load in the carcasses of the different mosquito populations (Fig 3.B); however, this variation was not significantly different for most comparisons, except between PET and POA populations (*p*-value = 0.01, Kruskal-Wallis test). It is interesting to note that this variation appears to have disappeared by 8 days post infection (Fig 3.D).

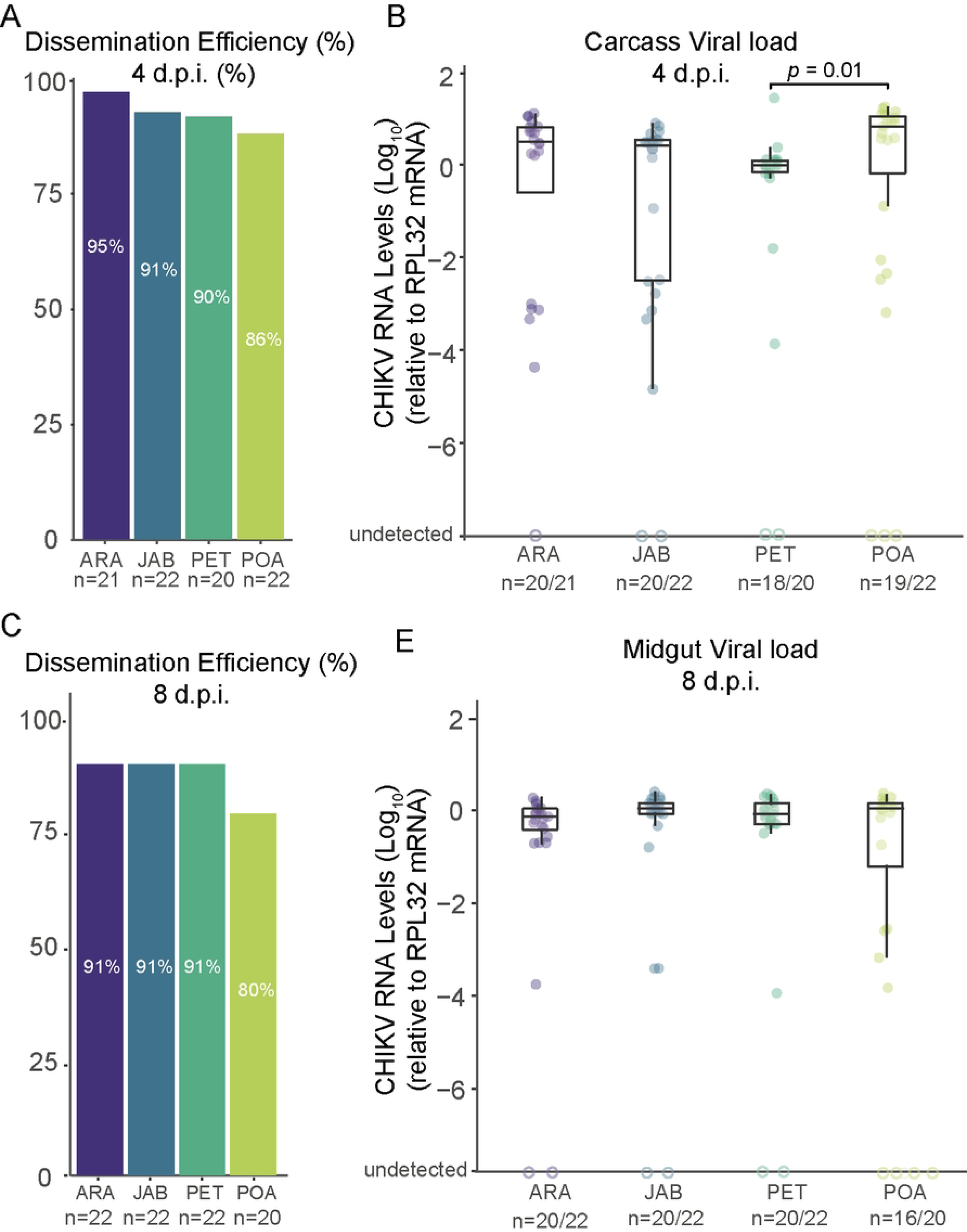

### High transmission efficiency of CHIKV across Brazilian *Aedes aegypti* populations

To assess the transmission efficiency of the CHIKV-ECSA strain we exposed mosquito females, that had an infectious blood meal 8 days previously, to AG129 naïve mice for a second blood meal. By exposing each female to one different AG129 mouse we were able to observe that most of the mice become infected with CHIKV (Fig 4.B). As shown in Fig 4.B, we found that transmission efficiency (the percentage of blood-fed mosquitoes that were able to infect the mouse out of the total number of blood-fed mosquitoes) varied from 80% in ARA population to 67% in PET population. Despite the observed variations, when we estimated the relationship between the transmission efficiencies and the different populations using a logistic regression model, we found no significant differences between the four mosquito populations (*p*-value = 0.9302, S3 Table).

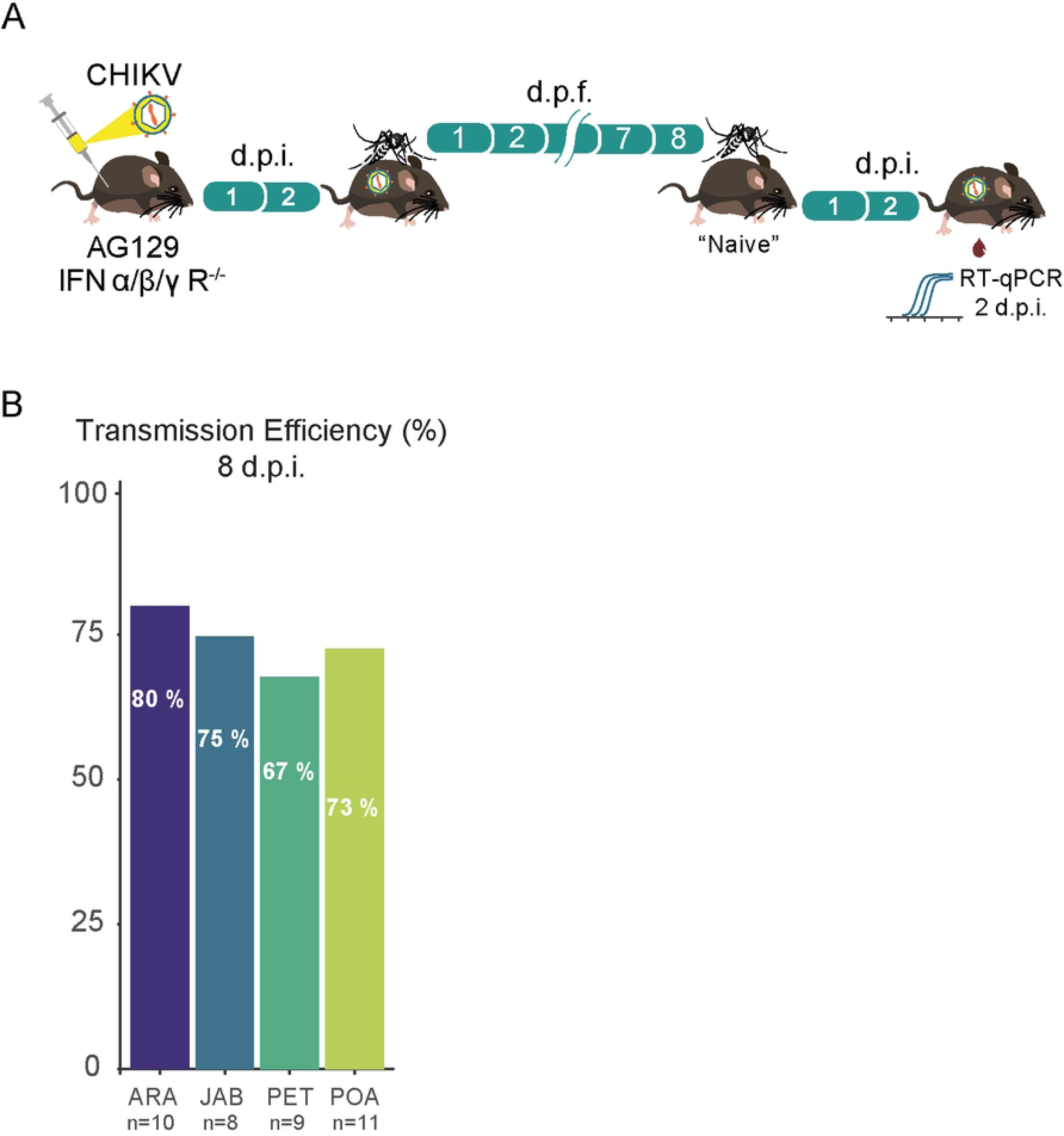

## Discussion

CHIKV has emerged as a global health threat over the past two decades, expanding from tropical and subtropical regions of Africa to Indian Ocean islands, the Indian subcontinent, South Asia, South America, and some regions in Europe [2,7]. Although two CHIKV lineages are confirmed to be circulating in Brazil: the Asian lineage and the ECSA lineage, surveillance studies indicate that the ECSA lineage is currently the pre-vailing genotype circulating in the country [10,26–28]. In this regard, evaluating the *Ae. aegypti* vector competence for the currently circulating CHIKV strain in this region is crucial to gain a better understanding of the role of the *Ae. aegypti* vector in CHIKV epidemiology in Brazil.

Using the viremic AG129 mice to infect the mosquitoes, our results confirm that the CHIKV-ECSA lineage exhibits a high level of infectivity across all Brazilian *Ae. aegypti* populations that were tested in this study, with infection rates reaching as high as 100% and no lower than 80% (Fig 2.B, D). Notably, the results also suggest that the CHIK-ECSA lineage is able to disseminate out of the midgut at a very early stage, since at just 4 days after the infectious blood meal, the majority of mosquitoes tested positive for the virus in their carcasses (Fig 3.A, B). Our results confirm previous findings that *Ae. aegypti* populations in South America exhibit high CHIKV dissemination efficiencies [29]. However, to our knowledge, this study is the first to report high infection rates and dissemination efficiencies using a locally isolated CHIKV strain that represents the currently circulating genotypes. Furthermore, by using AG129 mice to evaluate transmission competence that better resembles the natural cycle of transmission in nature, we have shown that the Brazilian populations are capable of efficiently transmitting the CHIKV-ECSA lineage, with transmission efficiencies ranging from 67% to 80%.

Overall, our results indicate that local *Ae. aegypti* populations are highly competent in being infected with and transmitting the CHIKV-ECSA strain circulating in Brazil. This suggests that Brazilian *Ae. aegypti* mosquitoes could play a significant role in disseminating this lineage across the country. These observations may help to explain why the CHIKV-ECSA lineage has experienced a significant expansion in Brazil in recent years, as well as the explosive emergence of CHIKV in Brazil and other South American countries. However, it is essential to note that the CHIKV Asian lineage is also present in Brazil, making it important to evaluate the susceptibility of *Ae. aegypti* populations to the local genotypes of this lineage. Additionally, other factors may also play a role in the epidemiology of CHIKV in Brazil. For example, high population densities in urban areas and a higher prevalence of *Ae. aegypti* over *Ae. albopictus* in these areas could also contribute to the transmission of the virus. Although *Ae. albopictus* has a more peri-urban distribution in Brazil, previous studies have reported that this species is a competent vector for different lineages of CHIKV [29]. Therefore, it is also crucial to assess the vector competence of local populations of this species for currently circulating CHIKV lineages in the South American regions. Taken together, our results may contribute to explain the recent epidemiology of CHIKV in Brazil and increased number of outbreaks associated with this ECSA lineage. Nonetheless, to gain a better understanding of the epidemiological dynamics of CHIKV, comprehensive studies are crucial, comparing currently circulating genotypes and including populations of both *Ae. aegypti* and *Ae. albopictus*. This information is vital for the development and implementation of effective strategies to control the spread of CHIKV and reduce the burden of this arbovirus on human populations.

## Acknowledgments

We are deeply thankful to João Marques, and their team members of RNAi Laboratory, UFMG, Brazil, for assistance with this project. We are grateful to Rivaldo Venâncio for the kindly provided CHIKV strain. We thank the Program for Technological Development of Tools for Health-PDTIS-FIOCRUZ for the use of its facilities. We are grateful to all members of the Mosquitos Vetores Group (MV-IRR/FIOCRUZ) and the World Mosquito Program Brazil. We are grateful for support from the CAPES finance code 001. L.A.M. and L.C.J.A. are fellows from CNPq. This work was also supported by grants from FAPEMIG (APQ-02760-17), CNPq, the Brazilian Ministry of Health (SVS-DEIDT and SCTIE-DECIT), the National Institutes of Health USA grant U01 AI151698 for the UWARN and INCT-EM.

